# Data-Independent Acquisition Mass Spectrometry as a Tool for Metaproteomics: Interlaboratory Comparison Using a Model Microbiome

**DOI:** 10.1101/2024.09.18.613707

**Authors:** Andrew T. Rajczewski, J. Alfredo Blakeley-Ruiz, Annaliese Meyer, Simina Vintila, Matthew R. McIlvin, Tim Van Den Bossche, Brian C. Searle, Timothy J. Griffin, Mak A. Saito, Manuel Kleiner, Pratik D. Jagtap

## Abstract

Mass spectrometry (MS)-based metaproteomics is used to identify and quantify proteins in microbiome samples, with the frequently used methodology being Data-Dependent Acquisition mass spectrometry (DDA-MS). However, DDA-MS is limited in its ability to reproducibly identify and quantify lower abundant peptides and proteins. To address DDA-MS deficiencies, proteomics researchers have started using Data-Independent Acquisition Mass Spectrometry (DIA-MS) for reproducible detection and quantification of peptides and proteins. We sought to evaluate the reproducibility and accuracy of DIA-MS metaproteomic measurements relative to DDA-MS using a mock community of known taxonomic composition. Artificial microbial communities of known composition were analyzed independently in three laboratories using DDA- and DIA-MS acquisition methods. DIA-MS yielded more protein and peptide identifications than DDA-MS in each laboratory. In addition, the protein and peptide identifications were more reproducible in all laboratories and provided an accurate quantification of proteins and taxonomic groups in the samples. We also identified some limitations of current DIA tools when applied to metaproteomic data, highlighting specific needs to improve DIA tools enabling analysis of metaproteomic datasets from complex microbiomes. Ultimately, DIA-MS represents a promising strategy for MS-based metaproteomics due to its large number of detected proteins and peptides, reproducibility, deep sequencing capabilities, and accurate quantitation.

## Introduction

Metaproteomics can provide direct functional readouts along with taxonomic information for a microbiome^1^, thereby holding great potential for medical^2^ and environmental^3^ microbiology applications. Metaproteomics uses bottom-up mass spectrometry-based proteomics to identify and quantify proteins in microbiome samples. The general shotgun proteomic process involves isolation of protein from samples, digestion into peptides, separation by liquid chromatography, and analysis of the peptides on the MS^4^. Over the past two decades^5–7^, the most common methodology for analysis of peptides has been Data-Dependent Acquisition mass spectrometry (DDA-MS).

In DDA-MS for shotgun proteomics, the mass spectrometer selects the most abundant ions that enter the instrument at any given time and isolates the ions for a controlled fragmentation^8^. Ideally, the mass spectrometer isolates a single peptide ion population per fragmentation spectrum, however that is not always the case^9^. A database search algorithm matches the experimental spectrum to *in silico* spectra generated from a protein sequence database to identify a peptide^10^. Previous studies have shown that this approach can effectively identify proteins in microbial communities at the species or sometimes even strain-level^11–13^, and more effectively measures percent biomass contributions of individual species than DNA sequencing-based methods^14^. For DDA-MS, the relative abundance of peptides at any given instant in the detector is partially stochastic due to ion interference and suppression^15^. As a result, DDA-MS is limited in its ability to reproducibly identify and quantify less abundant peptides and by extension proteins due to its inherent filtering for only the most abundant parent ions.

Advances in high resolution MS, faster scan speeds, and computational methods able to identify fragments from multiple peptides within a single spectrum^16,17^ resulted in the potential for an alternative acquisition method called Data-Independent Acquisition Mass Spectrometry (DIA-MS). DIA-MS represents a potential method for counteracting the deficiencies of DDA-MS. In DIA-MS, the mass spectrometer fragments all ions within a given range of mass-to-charge ratios, detecting the resulting ion fragments together, after which the instrument cycles to a new mass-to-charge range^18^. In DDA-MS, specific pairs of precursor and product ions are matched to peptides in a protein database. By contrast, in DIA-MS, there are two potential strategies. In the first, assorted precursors and products are combined into pseudo spectra which are searched against spectral libraries^19^, such as in the OpenSWATH suite^20^. These spectral libraries were originally constructed from DDA data, though they can now be generated directly from FASTA databases^21^ as in DIA-NN^22^ or the directDIA analysis of Spectronaut^23^. More recent approaches make use of chromatogram libraries, which make use of peptide spectra as well as retention time, as in EncyclopeDIA^24^. In the second method, peptides from the FASTA libraries are iteratively searched against the MS spectral data to ascertain their presence^25^ as in Group-DIA^26^, PECAN^27^, and the DIAmeter suite^28^. In both cases, DIA-MS theoretically allows for all ions regardless of their abundance to be detected, potentially allowing for more reproducible detection and quantification. Since the complexity of metaproteomics data precludes the efficient use of DDA-MS generated spectral libraries, it was the invention of directDIA software tools that allowed DIA to be feasible for metaproteomics. Several metaproteomics studies have been performed with DIA-MS, including analyses of human gut microbiota by Aakko et al.^29^ and Gomez-Varela et al.^30^ and examination of fermentation starter cultures by Zhao et al.^31^. Despite its use in the metaproteomics field, the accuracy of DIA-MS compared to DDA-MS is understudied. Zhao *et al*. validated the quantitative accuracy of DIA using a mock community from the perspective of log fold change^32^, but it did not evaluate quantitative accuracy from the perspective of exact protein input or attempt to evaluate identification accuracy using entrapment genomes of related species.

The objective of this study was to evaluate and compare the quality of identification and quantification of DDA-MS and DIA-MS metaproteomic approaches across three laboratories. We used a previously published 32-species mock community containing multiple bacteria, archaea, eukaryotes and viruses (Supplemental Table S1) to directly compare the ability of DIA-MS with DDA-MS to qualitatively and quantitatively characterize a microbiome^14^. We analyzed three types of mock communities in 4 biological replicates each: 1) using equal numbers of cells from each microorganism; 2) equal amounts of protein from each microorganism; and 3) uneven amounts of cells and protein from each organism (Figure 1A). To avoid that differences in sample processing influence the comparison, we prepared peptides in one laboratory and aliquoted them for subsequent analysis in the three participating laboratories. The 12 peptide samples were analyzed by the laboratories using DDA- and DIA-MS methods written for three different types of Orbitrap mass spectrometers. From there, the resulting DIA results were processed using directDIA analysis in Spectronaut. With the acquired data, we compared DIA-MS to DDA-MS from the following perspectives: a) the number of proteins and peptides that were detected in each sample using each LC-MS setup, b) the reproducibility of these protein and peptide measurements, c) the relative composition of the uneven samples, d) the per species depth of measurement, e) the relative quantitative accuracy for each species, and f) the number of proteins falsely identified when the protein database is constructed with species that are not present in the sample.

**Figure 1:**
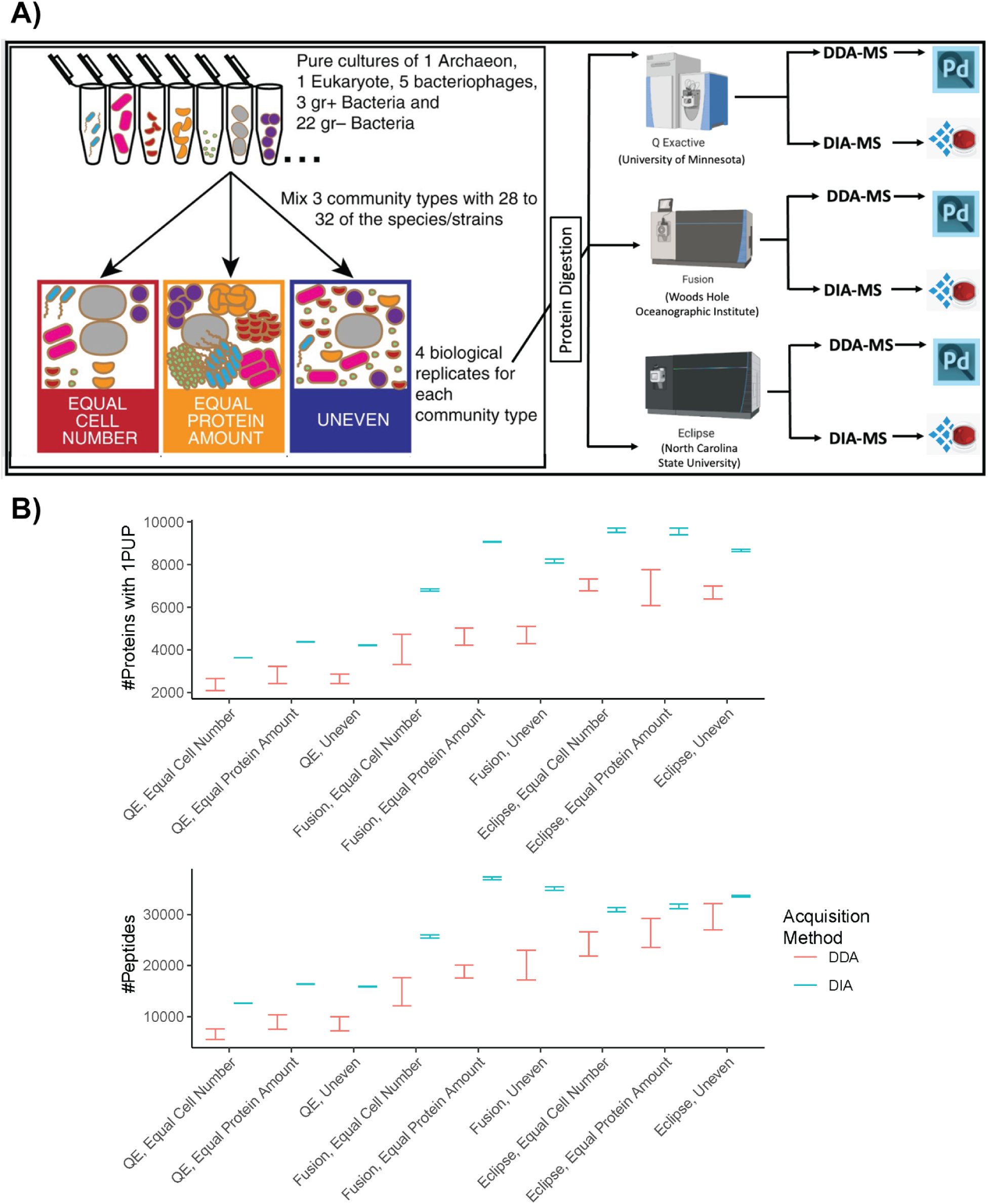
An outline of the generation of mass spectrometry data from the mock microbial communities. A) Illustration of mock community construction. 32 species and strains were used for the construction of three distinct community types. Four biological replicates each of the Equal Cell Number, Equal Protein Amount and Uneven were subjected to tryptic digestion. Peptides from all 12 samples were aliquoted, and sent to three laboratories for liquid chromatography and mass spectrometry. Figure adapted from Kleiner *et al.* 2017. DDA-MS data was analyzed using Proteome Discoverer (Pd) and the DIA-MS data was analyzed using Spectronaut software. B) Comparison of the number of proteins and peptides detected by DDA-MS versus DIA-MS using 95% confidence intervals (whiskers that do not overlap denote significance). Proteins were inferred by at least 1 protein unique peptide (PUP, a peptide that is unique to a specific protein accession in the database), and proteins and peptides were both inferred with an FDR of 1%.

## Methods

### Mock community sample preparation

Microbial mock communities (model microbiomes) were generated and frozen at −80°C for a previous study^14^. Briefly, 32 cultures of archaea, bacteria, eukarya and phages were used. We used four replicates, each of three of mock community types: 1) equal amounts of cells from each microorganism referred to as Equal Cell Number hereon); 2) equal amounts of protein from each microorganism (Equal Protein Amount hereon); and 3) randomized, uneven amounts of cells and protein from each organism (Uneven hereon). To generate the mock communities, cell pellets of each species were reconstituted and combined to generate multiple aliquots of each community and replicate before being snap frozen. Peptides from 12 samples (3 mock community types x 4 replicates) were generated using the filter-aided sample preparation (FASP) protocol described by Wiśniewski *et al*.^33^ The resulting peptides were desalted using Sep-Pak C18 Plus Light Cartridges (Waters, Milford, MA, USA) following the manufacturer’s instructions. Peptide concentrations were determined using the Pierce Micro BCA assay (Thermo Scientific Pierce, Rockford, IL, USA) following the manufacturer’s instructions. Aliquots of each sample were sent to the Griffin and Saito laboratories at the University of Minnesota and Woods Hole Oceanographic Institution, respectively, for analyses to complement analysis conducted at the Kleiner laboratory at NC State University.

### Construction of protein sequence databases

A protein sequence database for the mock community was generated by combining the proteomes of all species in the mock community into a protein database called “Mock_Comm_RefDB_V3_Clustered95.fasta” (112,580 sequences). Proteomes were acquired from UniProtKB or NCBI and are detailed in Supplemental Table S1. We made additional protein databases to test how the exclusion of specific species in the protein sequence database would affect the detection and quantification of other microbial proteins (Supplemental Table S1); a database called “Mock_Comm_RefDB_V3_Incomple1C95.fasta” (100,675 sequences) was generated that lacks the proteomes for *Rhizobium leguminosarum bv. viciae 3841*, *Pseudomonas denitrificans*, and *Pseudomonas fluorescens.* To further test this, the proteomes for *Pseudomonas pseudoalcaligenes*, *Salmonella enterica* Typhimurium LT2, and *Rhizobium leguminosarum bv. viciae VF39* were removed from Mock_Comm_RefDB_V3_Incomple1C95.fasta to generate the database “Mock_Comm_RefDB_V3_Incomple2C95.fasta” (84,216 sequences). To test the degree of misidentification of proteins that are not present in the mock community samples, a database was created by adding the protein sequences of *Bacteroides thetaiotaomicron* (different phylum), *Buttiauxella brennerae* (different genus), *Salmonella bongori* (different species), and *Tistrella mobilis* (different class) to Mock_Comm_RefDB_V3_Clustered95.fasta resulting in “Mock_Comm_RefDB_V3_Added1C95.fasta” (131,690 sequences). We also added these genomes from the Added 1 database to the Incomplete 1 and Incomplete 2 databases to generate“Mock_Comm_RefDB_V3_IncompleteAdded1C95.fasta” (119,785 sequences) and “Mock_Comm_RefDB_V3_IncompleteAdded2C95.fasta” (104,513 sequences) (Supplemental Table S1). Each protein database was clustered at 95% identity to remove redundant sequences using CD-HIT^34^.

### LC-MS/MS conditions

The microbiome samples were analyzed via LC-MS/MS in three separate laboratories using three separate instrument setups. For each mock community, single injections of the entire proteome were assayed in four biological replicate samples. In all laboratories, water with 0.1% formic acid was used as mobile phase A and acetonitrile with 0.1% formic acid was mobile phase B for LC applications.

For the Griffin lab at the University of Minnesota, samples were analyzed on a Thermo QExactive Quadrupole Orbitrap Hybrid Mass Spectrometer interfaced with an Ultimate 3000 UHPLC run in nano mode and plumbed with a nanoLC column packed with Luna C18 5µm resin (15 cm x 75 µm). For all LC-MS runs the same gradient was used, wherein the relative composition was held constant at 5% B for 5 minutes, followed by an increase to 35% B from 5 minutes to 95 minutes, an increase to 95% B from 95 to 100 minutes, a constant composition of 95% B from 100 to 110 minutes, a decrease from 95% B to 5% from 110 to 112 minutes, and a final re-equilibration stage held at 5% B from 112 minutes to 120 minutes. For DDA analyses, the instrument was run in positive mode using Full MS/dd-MS^2^ Top 15 mode. For the Full MS scan the resolution was 35,000 with an automatic gain control (AGC) target of 1e6, a maximum injection time (IT) of 30 milliseconds, and a scan range of 400-1600 m/z. Data-dependent MS^2^ were collected at a resolution of 17,500 with an AGC target of 1e6, a maximum IT of 50 milliseconds, an isolation window of 2.0 m/z and a scan range of 200 - 2000 m/z. To conduct DIA analyses, Full MS scans were combined with DIA scans, both of which were run in positive mode. For the Full MS scan, the resolution was 35,000 with an AGC target of 1e6, a maximum IT of 200 milliseconds, and a scan range of 385-1015 m/z. For the DIA scan, the resolution was set to 17,500 with an AGC target of 1e6, a loop count of 25, an isolation window of 24 m/z, and two sets of staggered DIA scan windows from 400 to 1000 m/z and from 388 to 988 m/z.

In the Saito lab at Woods Hole Oceanographic Institution, measurements were performed on a Thermo Fusion Orbitrap Tribrid Mass Spectrometer interfaced with an Ultimate 3000 RSLCnano system plumbed with a nanoLC column packed with C18 Reprosil-Gold 3µm resin (15 cm x 100 µm). For all LC-MS runs a gradient was used which began with 2% B for the first 10 minutes, after which the % B increased from 2 to 28% from 10 to 90 minutes, followed by an increase of 28 to 35% B from 90 to 102 minutes, 35 to 95% B from 102 to 108 minutes, a constant 95% wash from 108 to 109 minutes, a decrease of 95 to 2% B from 109 to 110 minutes, culminating with a 2% re-equilibration from 110 to 120 minutes. For DDA analyses the instrument was run in positive mode with a full scan at resolution 240,000, a scan range of 380-1280 m/z, standard AGC targeting, an automatic maximum injection time mode, an intensity threshold of 1.0e3, and cycle time mode with 2 seconds between full scans; following the full scan was a data-dependent MS^2^ scan in which the normalized collision energy was 27, the ion trap was employed as the ms2 detector. In DIA experiments there were four scans; first a master scan ran in positive mode at resolution 60000 with a scan range of 385-1015 m/z, next a DIA experiment in positive mode with 25 scan windows of 24 m/z width and a range of 400-1000 m/z at 30,000 resolution and normalized collision energy of 27, followed by a second master scan, and concluding with a second DIA experiment in positive mode with 25 scan windows of 24 m/z width and a range of 412-988 m/z.

In the Kleiner lab at North Carolina State University, peptides were separated along a 140 minute reverse phase gradient using an Ultimate 3000 RSLCnano system interfaced with a 75 cm x 75 µm analytical EASY-Spray PepMap RSLC C18 column. The gradient was as follows: 5% to 31% B for 102 minutes, 31% B to 50% B for 18 minutes, and 50% to 99% B for 20 minutes. This system was coupled to a Thermo Orbitrap Eclipse Tribrid Mass Spectrometer. For DDA analysis the instrument was run in positive mode with a full scan at resolution 60,000, a scan range of 380-1600 m/z, and AGC target of 300% (3.0 x 10^6^ charges) a maximum injection time of 200 ms. For data-dependent MS^2^ acquisition, there were 15 dependent scans, at a minimum intensity threshold of 5×10^3^, with a 25 minute exclusion list. For MS^2^ scans ions were fragmented with an HCD collision energy of 27% and measured at a resolution of 15,000, an AGC target of 100% (1 x 10^5^ charges) and a maximum injection time of 50 ms. For the DIA experiments there was first a full scan run in positive mode at a resolution of 60,000, with a scan range of 380-1600 m/z, a 200 ms maximum injection time, and an AGC target of 300%. Data-independent MS2 scans were collected on a 30 spectra loop in isolation windows of 10 m/z over the range of 384-1000 m/z. MS^2^ scans were fragmented at an HCD collision energy of 27% and measured at a resolution of 15,000, an AGC target of 100%, and 50 ms maximum injection time.

### Data Analysis

Raw DDA mass spectrometry files were searched against microbial community protein sequence FASTA files using Proteome Discoverer (v2.3)^35^. The processing steps used Sequest HT and Percolator to match peptide spectra to the mock community protein sequence FASTA files. In Sequest HT, trypsin was selected as the enzyme with a maximum missed cleavage number of 2. The precursor tolerance was set to 10 ppm and the fragment tolerance was set to 0.1 Da. For spectrum matching, b- and y-ions were selected. Methionine oxidation, deamidation (N,Q,R), and protein N-terminal acetylation were set as dynamic modifications while carbamidomethylation of cysteine was set as a fixed modification. Each raw file was searched separately and consensus was only used to output the data from each individual search.

The raw mass spectrometry files were imported into the Spectronaut software^23^ version 18.3.230830 along with the protein sequence FASTA databases. Spectronaut was then run with the directDIA+ workflow with 10 ppm MS1 and MS2 relative tolerance at the Calibration search and Main Search levels and a false discovery rate of 0.01 at the PSM, peptide, and protein group level. Trypsin/P was selected as the protease with two missed cleavages allowed and cysteine carbamidomethylation was set as a fixed modification. In addition, protein N-terminal acetylation, glutamine/arginine/asparagine deamidation, and methionine oxidation were selected as variable modifications to peptides.

Proteins were inferred if they had at least 1 protein unique peptide and a protein FDR <1%. The mean number of protein groups and peptides detected across each method were compared to one another using 95% confidence intervals. The degree of detected peptide overlap between replicates was determined using UpSet plots. Comparison of measured and reference percent abundances of constituent species in Uneven samples was conducted qualitatively using a bubble plot and quantitatively via Spearman correlation. Quantitative comparison of DDA- and DIA-MS runs was done using the metric below comparing the measured percent abundances to the theoretical percent abundances in the constant protein and uneven protein samples.

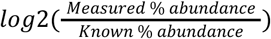

The 95% confidence intervals for this metric were calculated using the values from each species present in the sample and were compared with one another. We calculated the 95% confidence intervals of the mean total number of protein matches for alternative protein sequence databases to examine the effect of additional or missing protein sequences on the number of protein matches. To determine the rate of false positive detections, the 95% confidence intervals were calculated for the mean percentage of protein identifications to added and incomplete databases described previously.

Figures were generated using the R statistical computing software v4.2.2 with the Rmisc^36^ and ggplot2^37^ packages.

## Results

### DIA methods result in more proteins and peptides detected than with DDA regardless of protein inference method

To compare the number of identifications generated by our DIA approaches relative to our DDA approaches we compared the 95% confidence intervals of the mean between DIA and DDA for all measurements and community types (Figure 1B). DIA consistently detected significantly more proteins using 1 protein-unique peptide (PUP) for protein level inference and peptides than DDA methods at a protein false discovery rate (FDR) of <1% across all communities (Equal Cell Number, Equal Protein, and Uneven) and instrument types (Supplemental Table S2). This observation held true using other protein inference methods as well (minimum of 2 PUPs, parsimony-based protein grouping, etc.).

### DIA methods are more reproducible in their detection of peptides

To compare measurement reproducibility between DIA and DDA, we analyzed how many peptides were reproducibly detected across all the measurements using UpSet plots (Figure 2; Supplemental Figure S1). Using DIA methods regardless of method and community type almost all the peptides detected were detected in all four replicates. In contrast, across all community types the DDA methods had a large number of peptides detected in only one of the four replicates. These results show that DIA methods were more reproducible than DDA methods.

**Figure 2:**
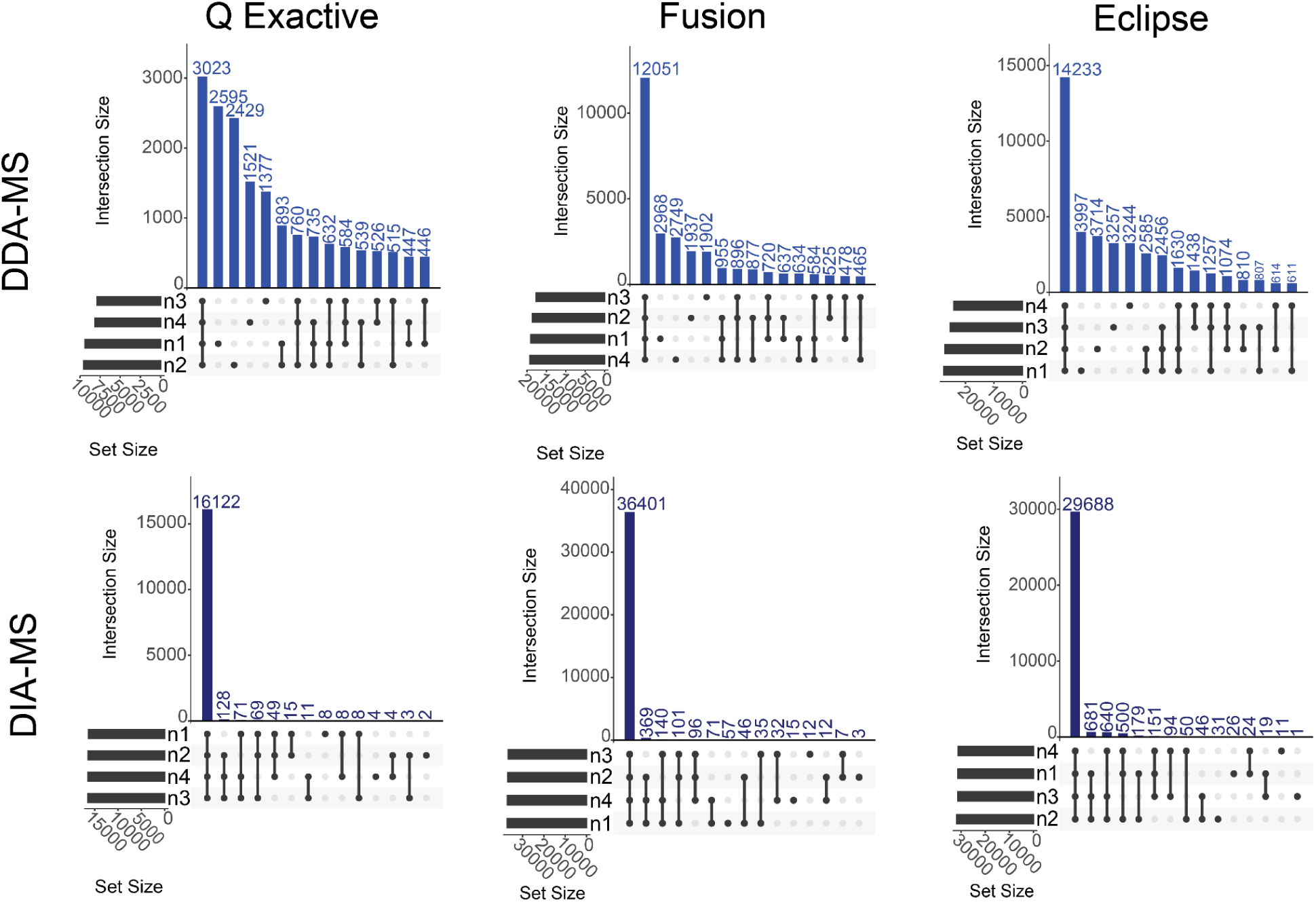
Data Independent Analysis mass spectrometry yields more peptide identifications more reproducibly. UpSet plots of peptide identification reproducibility of Uneven communities across three mass spectrometers (QExactive, Fusion, and Eclipse) under DDA- and DIA-MS analysis. The variables n1 through n4 represent MS runs of the four replicates. Peptides were filtered at 1% FDR.

### DIA methods result in comparable detection of proteins from low abundance species to DDA

Since the compositions of the mock microbial communities used in this study were known, we were able to determine if our methods accurately reproduced the known species abundances in terms of their proteinaceous biomass contributions. We compared the percent abundance of the species using our DDA and DIA methods to the known amount of protein added for each species to the community (Figure 3A). In all analyses, DDA and DIA both detected similar percent compositions to the known community composition, with Spearman correlation values between the known and measured percent abundances for DDA- and DIA-MS data averaging at 0.946 and 0.924, respectively (Supplemental Figure S2). In all instances, we detected S*almonella enterica* Serovar Typhimurium LT2 (LT2) as underrepresented relative to the known percent abundance, while *Cupriavidus metallireducens* (Cup) and *Escherichia coli* (K12) had inflated percent abundances relative to the known composition.

**Figure 3:**
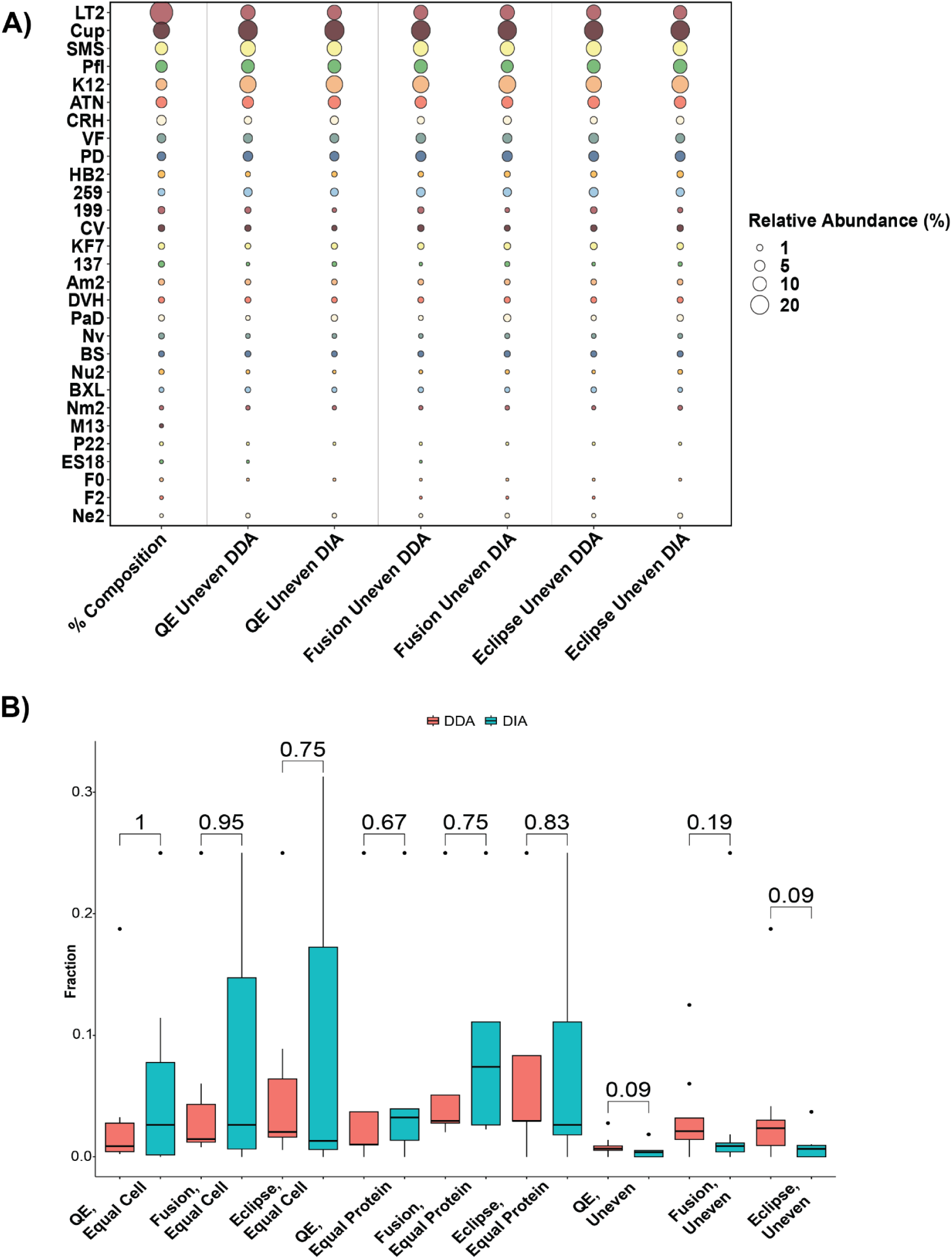
DIA-MS recapitulates the proportions of the microbial communities seen in DDA-MS. a) Known community composition of the Uneven mock community samples compared to the percent abundance of each species as determined by multiple instruments, acquisition methods and software types. Percent abundances were calculated based on the summed spectral counts (DDA-MS) or summed intensities (DIA-MS) of each species divided by the total MS signal^14^ b) Number of detected proteins divided by the total number of protein-coding genes in the genome of bottom 25% least abundant species in microbiome samples. DDA- and DIA-MS samples were compared using a Mann-Whitney-Wilcoxon test, with the p-values displayed above the bars. Boxes represent the 1st through 3rd quartile of the measured values of multiple measured species, while whiskers extend up to 1.5 times the interquartile range from the box to encapsulate data points outside the interquartile range; data points beyond this range are expressed as points on the graph.

By examining the less abundant species in the mock communities, we investigated whether DDA-MS or DIA-MS was able to detect a greater fraction of the total proteome (the number of detected proteins out of the total number of protein-encoding genes) of these species. We found that we detected more peptides from the low abundance species in DDA relative to DIA in the Uneven samples (Supplemental Table S2). To further investigate this, we quantified the fraction of the total proteome for the 25% least abundant species based on sample formulation (Figure 3B and Supplemental Table S1). Interestingly, we found that DDA-MS trended towards slightly higher fractions of low abundance species in the Uneven samples, but this was not significant by the Mann–Whitney–Wilcoxon test (Figure 3B). Thus while DIA outperformed DDA for the whole community analysis (all species in mock communities, Figures 1 and 2), this advantage disappears when focused solely on rarer organisms, and both methods are equally adept at detecting low abundance species, though DDA detects more peptides. In examining the individual species, DIA-MS tended to detect higher fractions of the more abundant species than DDA-MS (Supplemental Table S3).

### DIA methods result in a comparable level of quantitative accuracy to DDA

To assess the quantitative accuracy of DIA methods for proteomics, we calculated the log_2_ of the measured percent abundance of each species in the community divided by the expected percent abundance of that species for each method and community type and then calculated the 95% confidence intervals of the mean value (Figure 4a). In this analysis, values of zero mean that the measured percent abundance of a species is equal to its expected abundance based on the physical amount of that organism’s protein that was input into the mock community. For all methods and community types the confidence intervals overlapped between DDA and DIA indicating that they do not differ in accuracy. In addition, for some of the methods the confidence interval overlapped with zero in the equal cell and uneven communities suggesting that the quantitative accuracy was within a confidence interval of 95% for these measurements.

**Figure 4:**
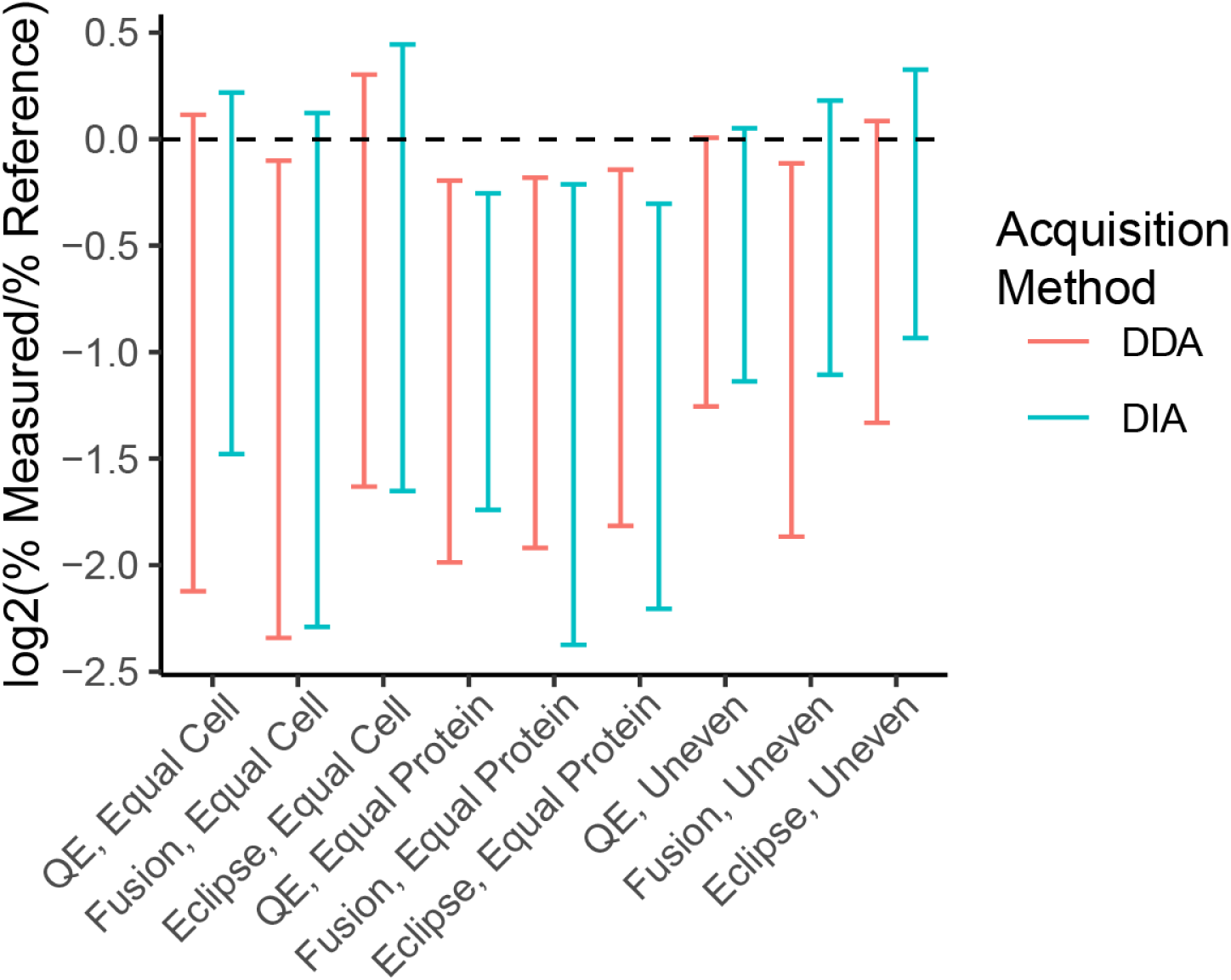
Assessment of the quantitative accuracy of the mass spectrometry methods for determining species abundances in the mock communities. Data represent the 95% confidence intervals of the base 2 log of the ratio of the measured species abundances to the known species abundances.

### DIA and DDA methods result in a comparable number of false protein identifications when challenged with incomplete databases with entrapment organisms

There is a degree of unavoidable uncertainty when it comes to protein sequence database design for experimental metaproteomics datasets. Depending on the methods used for database generation, protein sequences from microorganisms present in the sample are potentially missing in the search database and conversely protein sequences from microorganisms that are not present in the sample might get included in the search database^38^. The defined nature of this sample set allowed us to investigate the effects of including protein sequences from species in the database that are not present in the samples and excluding protein sequences from species present in the sample from the database. To do this, we generated five additional databases: incomplete 1 (I1), incomplete 2 (I2), added (A1), incomplete added 1 (A1I1), and incomplete added 2 (A1I2) (Supplemental Table S2). Incomplete databases progressively increase the number of genomes missing. For incomplete 1, *Rhizobium leguminosarum* bv. *viciae* 3841, *Pseudomonas denitrificans*, and *Pseudomonas fluorescens* were removed. For incomplete 2, *Pseudomonas pseudoalcaligenes*, *Salmonella enterica* Typhimurium LT2, and *Rhizobium leguminosarum* bv. *viciae* VF39 were also removed. For all three added databases, the protein sequences of *Bacteroides thetaiotaomicron*, *Buttiauxella brennerae*, *Salmonella bongori*, and *Tistrella mobilis* were added as entrapment sequences to generate potential false protein identifications. We selected these four bacterial genomes because they had varying amounts of genetic distance from the organisms present in the mock community: *B. thetaiotaomicron* was in a different phylum from any of the bacteria in the mock community, *T. mobilis* was in a different class from any bacteria in the mock community, *B. brennerae* shared the same family with multiple members of the community but was in a different genus, and *S. bongori* shared the same genus but is a different species from *S. enterica* which was removed in the incomplete 2 database. We hypothesized that we would identify some number of false protein identifications from all the added species, but that we would detect more false protein identifications in the incomplete added 2 database due to peptides being assigned to *S. bongori* due to the removal of *S. enterica* from the incomplete 2 database. We use the term “false protein identifications” in a loose sense here, as it is likely that cross-species identification of peptides and proteins occurs where the correct peptides and proteins are identified with sequences of a closely related species in the absence of the sequences from the correct species if the sequences share stretches in which they are identical.

We found that regardless of the inference method, removing protein sequences from the database had a greater impact on the total number of identifications than adding entrapment protein sequences (Figure 5A-5F). For all methods, the confidence intervals generally overlapped between the standard, added 1, incomplete 1, and incomplete added 1 databases; however, the number of identifications significantly decreased when the incomplete 2, and incomplete added 2 databases were used.

**Figure 5:**
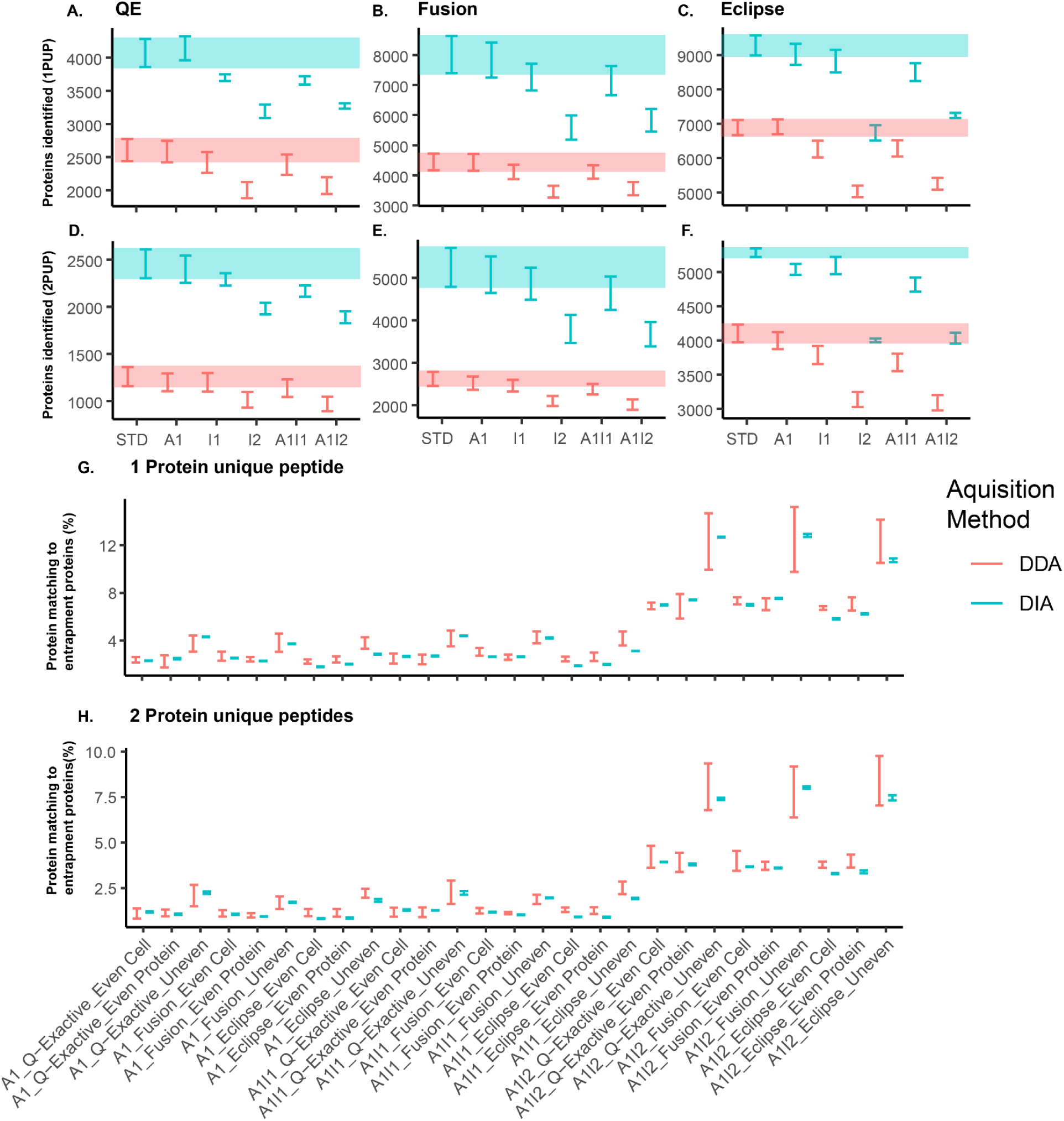
DIA measurements misidentify proteins at an equal or lower rate than DDA. The 95% confidence intervals for number of proteins identified using at least one protein unique peptide (1PUP) for each of the different database configurations Added (A1), Incomplete Added 1 (A1I1), Incomplete Added 2 (A1I2), Incomplete 1 (I1), Incomplete 2 (I2), and the standard database (STD) for the (A) QE, (B) Fusion, and (C) Eclipse measurements. The 95% confidence intervals for the number of proteins identified using at least 2 protein unique peptides (2PUP) for each of the different database configurations for the (D) QE, (E) Fusion, and (F) Eclipse measurements. Shaded bands represent the 95% confidence interval of the STD database. (G) The 95% confidence intervals for the percentage of the 1PUP proteomes that matched entrapment protein sequences in the A1, A1I1, and A1I2 databases. (H) The 95% confidence interval for the percentage of the 2PUP proteomes that matched entrapment protein sequences in the A1, A1I1, and A1I2 databases. Protein FDR was controlled at 1%.

With the added databases we investigated whether the addition of entrapment protein sequences generated false protein identifications in DIA-MS and DDA-MS (Figure 5G-5H). Across all methods and community types, we observed that DIA and DDA methods had similar rates of false protein identifications. However, the rate of false protein identifications decreased when we used a stricter protein inference criterion (2 protein unique peptides). For the incomplete added 2 database the removal of sequences from the genome of *Salmonella enterica* Typhimurium LT2 led to a substantial increase in the detection of proteins from *S. bongori*, which led to an increase in the rate of false protein identifications most likely due to *S. enterica* peptides being assigned to the homologous sequences of *S. bongori*. That being said, many proteins were also detected from *B. brennerae* and less from T. *mobilis* and *B. thetaiotaomicron,* which are likely could still be matches too similar peptides and homologous proteins, but could also be true false positive identifications (Supplemental Figure S3).

## Discussion

Our goal for this study was to compare the depth, quantitative accuracy, and identification accuracy of DIA-MS methods for metaproteomics to DDA-MS methods for metaproteomics using three different mock microbial communities of known composition. Ultimately, the comparisons between DDA- and DIA-MS were made within each lab and broad inferences can be made collectively. Similar to previous studies, we found that DIA-methods in general had more protein and peptide identifications than DDA methods across all MS platforms^32^. We also observed that the peptide identifications using DIA methods were substantially more reproducible than DDA methods. For DDA-MS, we found that some peptides (∼30%) were identified across all four replicates regardless of method or community type, but the majority of peptides were found in only one of the four replicates on all three instruments (Figure 2 top panels). In contrast, for DIA-MS we observed that the majority of peptides were identified in all four replicates across all MS platforms (Figure 2 bottom panels). This can likely be explained by the stochastic nature of ion fragmentation in DDA-MS methods, which results in a lack of reproducibility between runs^39^. In contrast, DIA-MS^40^ has increased amounts of reproducibly identified peptides due to the non-stochastic nature of peptide detection in the DIA-MS analyses as compared to the DDA-MS analyses. In the recent Critical Assessment of MetaProteome Investigation (CAMPI) study^41^, which only used DDA-MS data, different wet-lab workflows showed considerable overlap in protein subgroups, particularly among the most abundant ones. However, the peptide overlap across these workflows was rather limited. Our study shows that DIA-MS for metaproteomics is substantially more reproducible at the peptide level than DDA-MS, providing the potential to increase the reproducibility between experiments and therefore confidence in metaproteomic analysis.

We further investigated the quantitative accuracy of DIA methods relative to DDA methods by comparing the relative measured abundances of the species that made up the mock communities to the species’ known relative abundances. We found that the DIA methods had comparable accuracy to the DDA methods, which had already been shown to be more accurate than sequencing-based methods for the quantification of proteinaceous biomass using the same mock communities^14^. Our results are in line with a previous study that showed that DIA had a comparable quantitative accuracy to DDA methods using a 12 member synthetic community^32^; the authors only presented quantitative accuracy with regards to log fold changes. In this study we show that DDA and DIA methods are comparable for determining the percent abundance of an organism within a mock community across multiple domains of life and for a much larger number of organisms.

Due to the chimeric nature of MS2 spectra, it was possible that DIA-MS metaproteomic methods would have a greater rate of false positive identifications than DDA-MS despite producing a deeper proteome. With directDIA identification methods, DIA-MS metaproteomics is just as dependent as DDA-MS on the quality of the protein database used for identification of peptides and proteins^38^. To investigate whether DIA-MS was more prone to false identifications than DDA-MS, we created additional databases that were missing the proteins from select species known to be in the community and also added proteins from four species known to not be in the sample and of varying phylogenetic distance from the members of the community. In general, DIA and DDA methods identified a similar percentage of proteins from species known to not be in the sample relative to the total number of identifications. We also found that the combined effect of removing genomes known to be in the sample and adding genomes known to not be in the sample had a significant effect on the number of identifications, and that this particular combination had a substantial effect on identification accuracy by increasing the number of proteins from species known to not be in the sample to >7% of the identifications, even with the stricter inference threshold of at least 2 protein unique peptides. The majority of the “false positives” in these data across databases and MS platforms belonged to *Salmonella bongori*, a species with approximately 83.6% sequence identity^42^ to the *S. enterica* serovar typhimurium strains known to be present in the original mock community samples analyzed in this study and specifically removed from two of the databases. We acknowledge that this analysis is not a true measure of the false positive rate since we did not add an equal number of entrapment proteins as the total size of the protein database, as such many of the misidentified proteins were due to peptide matches belonging to homologous proteins from a closely related species removed from the database or were due to peptide matches that should have instead been to a similar peptide from a conserved region within a homologous protein that actually was in the sample. Despite this, we think that this analysis is useful because it shows that DDA-MS and DIA-MS have similar false identification rates and highlights the importance of careful database design for metaproteomic studies, regardless of whether a DDA or DIA method is used because having a database that is not comprehensive and has protein sequences that do not belong can lead to less identifications and a high number of misidentified proteins overall.

In our work we identified several limitations of bioinformatic analysis of DIA metaproteomics datasets, which need to be addressed in the future to make it a widely usable approach. We tried multiple open-source softwares, but were ultimately only able to process our data with Spectronaut. For Spectronaut, which we ended up using for DIA data analysis, it depended on the version number if data processing could be completed in a reasonable time frame. For example, while version 17 of Spectronaut was able to complete processing of individual samples in the direct DIA+ mode within a few hours for each sample, version 18 ran for over 15 hours on an individual sample. Second, the protein sequence databases that we used for protein identification in this study were relatively small (112,592 protein sequences), as compared to databases used for metaproteomics of more complex, real life samples such as fecal material or soil samples (>500,000 to millions of protein sequences)^13,43^. While DIA metaproteomics is superior for low complexity samples that only require small protein sequence databases, further testing and software development will be needed to make DIA metaproteomics feasible and accessible for more complex samples. A focus on faster run times, the capability to parallelize on multiple servers, and specific handling for large databases would go a long way towards increasing the useability of DIA for metaproteomics. To our knowledge all DIA metaproteomic studies to date done on more complex samples have had to reduce their database size to enable data analysis^30,44^.

In this work we examined the efficacy of DIA- relative to DDA-MS for metaproteomics across multiple LC-MS/MS setups and software suites. Ultimately we found that DIA-MS was able to identify more proteins than DDA-MS and had much more reproducible peptide and protein identifications across replicate measurements. Furthermore, DIA demonstrated accurate quantification in uneven, constant protein, and constant cell samples. Yet while DIA-MS identified comparable levels of low-abundance species (bottom quartile) per mock community sample, it did not outperform DDA as it did in peptide and protein identifications when all of the community was included. In particular, the two tribrid instruments showed trends for DDA being more sensitive than DIA (Figure 3B) that may be due to the combination of sensitivity of the ion trap and small DDA ms2 fractionation windows, whereas the combination of relatively large isolation windows used (10-24m/z) and low precursor intensities found in the low abundance species could make obtaining sufficient ions for high quality ms2 scans more challenging in DIA. Finally, when entrapment protein sequences are included in the FASTA database, DDA-MS and DIA-MS have similar levels of false positive detection. Taken together, our results suggest that DIA-MS has the potential for superior performance for metaproteomics analyses as compared to DDA-MS. The high-quality MS datasets generated in our experiments are available for the metaproteomics community to explore for testing and the development of new algorithms and methodologies. While this superior performance is already available for low complexity microbial communities for which relatively small protein sequence databases (∼100,000 sequences) are needed,^11,45^, future work in the optimization of search algorithms is needed to make DIA metaproteomics feasible for more complex microbial communities that require use of larger protein sequence databases which have millions of sequences. In addition, future work will examine the capabilities of other cutting edge platforms for DIA in metaproteomics such as DIA-PASEF on the timsTOF^46^, ZenoSWATH^47^, and the Orbitrap Astral^48^.

## Associated Data

The raw data, search results, and metadata SDRF file^49^ generated with lesSDRF^50^ can be found in the PRIDE database^51^ under the entry PXD054415 [reviewer account name, password LWvi2Ps4Ws2k]. Search results from alternate protein sequence databases as well as all protein sequence databases used can be found at Zenodo https://zenodo.org/doi/10.5281/zenodo.13376413.

## Acknowledgements

This work has benefited from collaborations facilitated by the Metaproteomics Initiative (https://metaproteomics.org/) whose goals are to promote, improve and standardize metaproteomics^52^. Part of the LC-MS/MS measurements were made in the Molecular Education, Technology, and Research Innovation Center (METRIC) at North Carolina State University. This work was funded by a postdoctoral fellowship through the National Institutes of Health grant 2T32DK007737-26 (JABR), the National Institute Of General Medical Sciences of the National Institutes of Health under Award Numbers R35GM138362 (MK) and R01GM135709 (MAS), National Science Foundation OCE-2123055 (MAS) and the U.S. Department of Agriculture National Institute of Food and Agriculture under award No. 2021-67013-34537 (MK). T.V.D.B. acknowledges funding from the Research Foundation Flanders (FWO) [1286824N]. We would like to thank Biognosys for offering license to perform Spectronaut searches. Authors disclose that there are no conflicts of interest.

