## Supplemental Information for "Data-Independent Acquisition Mass Spectrometry as a Tool for Metaproteomics: Interlaboratory Comparison Using a Model Microbiome"

### Supplemental Information for Data-Independent Acquisition Mass Spectrometry as a Tool for Metaproteomics: Cross-Laboratory Methodological Comparisons Using a Model Microbiome

Rajczewski, A. T.[1], Blakeley-Ruiz, J. A.[2], Meyer, A.[3], Vintila, S.[2], McIlvin, M.R.[4], Van Den Bossche, T.[5, 6], Searle, B.C.[7], Griffin, T.J.[1], Saito, M.[4], Kleiner, M.[2], Jagtap, P.D.[1]

[1] Department of Biochemistry, Molecular Biology, and Biophysics, University of Minnesota, Minneapolis MN USA
[2] Department of Plant and Microbial Biology, North Carolina State University, Raleigh NC USA
[3] MIT-WHOI Joint Program in Oceanography/Applied Ocean Science and Engineering, Department of Chemistry, Woods Hole Oceanographic Institution, Woods Hole MA USA, Department of Earth, Atmospheric, and Planetary Sciences, Massachusetts Institute of Technology, Cambridge MA USA
[4] Department of Marine Chemistry and Geochemistry, Woods Hole Oceanographic Institution, Woods Hole MA USA
[5] VIB-UGent Center for Medical Biotechnology, VIB,, Ghent Belgium
[6] Department of Biomolecular Medicine, Ghent University, Ghent Belgium
[7] Department of Chemistry and Biochemistry, The Ohio State University, Columbus OH USA

Supplemental Table S1: Protein sequence database components used in the proteomic analyses of composite microbiome samples

| **Label** | **Species** | Source of protein sequences | Original Database | Incomplete 1 | Incomplete 2 | Added1 | Added Incomplete1 | Added Incomplete2 |
|---|---|---|---|---|---|---|---|---|
| NV | *Nitrososphaera viennensis* | GCA_000698785.1 | Y | Y | Y | Y | Y | Y |
| 841 | *Rhizobium leguminosarum bv. viciae* 3841 | UP000006575 | Y | N | N | Y | N | N |
| PaD | *Paracoccus denitrificans* | Used RAST. Available in the protein sequence database on PRIDE. | Y | Y | Y | Y | Y | Y |
| Cup | *Cupriavidus metallidurans* | UP000002429 | Y | Y | Y | Y | Y | Y |
| CV | *Chromobacterium violaceum* | Used RAST. Available in the protein sequence database on PRIDE. | Y | Y | Y | Y | Y | Y |
| Nu1 | *Nitrosomonas ureae* | UP000056699 | Y | Y | Y | Y | Y | Y |
| DVH | *Desulfovibrio vulgaris* | UP000002194 | Y | Y | Y | Y | Y | Y |
| K12 | *Escherichia coli* | UP000000625 | Y | Y | Y | Y | Y | Y |
| PD | *Pseudomonas denitrificans* | UP000012082 | Y | N | N | Y | N | N |
| KF7 | *Pseudomonas pseudoalcaligenes* | GCA_000262065.3 | Y | Y | N | Y | Y | N |
| Pfl | *Pseudomonas fluorescens* | 2617270901 (IMG) | Y | N | N | Y | N | N |
| HB2 | *Thermus Thermophilus* | UP000000592 | Y | Y | Y | Y | Y | Y |
| CRH | *Chlamydomonas reinhardtii* | GCF_000002595.1 | Y | Y | Y | Y | Y | Y |
| LT2 | *Salmonella enterica Typhimurium* (3 strains combined) | UP000001014 | Y | Y | N | Y | Y | N |

| | | | | | | | | |
|---|---|---|---|---|---|---|---|---|
| ATN | *Agrobacterium tumefaciens* | UP00000813 | Y | Y | Y | Y | Y | Y |
| VF | *Rhizobium leguminosarum bv. viciae* VF39 | GCA_000427765.1 | Y | Y | N | Y | Y | N |
| AK199 | *Roseobacter sp.* AK199 | Used RAST. Available in the protein sequence database on PRIDE. | Y | Y | Y | Y | Y | Y |
| BXL | *Burkholderia xenovorans* | UP000001817 | Y | Y | Y | Y | Y | Y |
| Ne1 | *Nitrosomonas europaeae* | UP000001416 | Y | Y | Y | Y | Y | Y |
| Nm1 | *Nitrosospira multiformis* | UP000002718 | Y | Y | Y | Y | Y | Y |
| Am2 | *Alteromonas macleodii* | UP000006296 | Y | Y | Y | Y | Y | Y |
| SMS | *Stenotrophomonas maltophilia* | GCF_000613205.1 | Y | Y | Y | Y | Y | Y |
| BS | *Bacillus subtilis* | UP000001570 | Y | Y | Y | Y | Y | Y |
| 137 | *Staphylococcus aureus* ATCC 13709 | Used RAST.. Available in the protein sequence database on PRIDE | Y | Y | Y | Y | Y | Y |
| 259 | *Staphylococcus aureus* ATCC 25923 | GCA_000756205.1 | Y | Y | Y | Y | Y | Y |
| ES18 | Phage ES18 | UP000000970 | Y | Y | Y | Y | Y | Y |
| F0 | Phage F0 | UP000009070 | Y | Y | Y | Y | Y | Y |
| F2 | Phage F2 | UP000002127 | Y | Y | Y | Y | Y | Y |
| M13 | Phage M13 | UP000002111 | Y | Y | Y | Y | Y | Y |
| P22 | Phage P22 | UP000007960 | Y | Y | Y | Y | Y | Y |
| BT | *Bacteroides thetaiotaomicron* | UP00000141 | N | N | N | Y | Y | Y |
| BBE | *Buttiauxella brennerae* | UP00007841 | N | N | N | Y | Y | Y |
| SBI | *Salmonella bongori* | UP00027208 | N | N | N | Y | Y | Y |
| TMO | *Tistrella mobilis* | UP00000525 | N | N | N | Y | Y | Y |

Supplemental Figure S1: UpSet plots of peptide reproducibility of Equal Cell Number (red) and Equal Protein Amount (green) communities across three mass spectrometers (QExactive, Fusion, and Eclipse) under DDA- and DIA-MS analysis. The variables n1 through n4 represent MS runs of the four replicates.

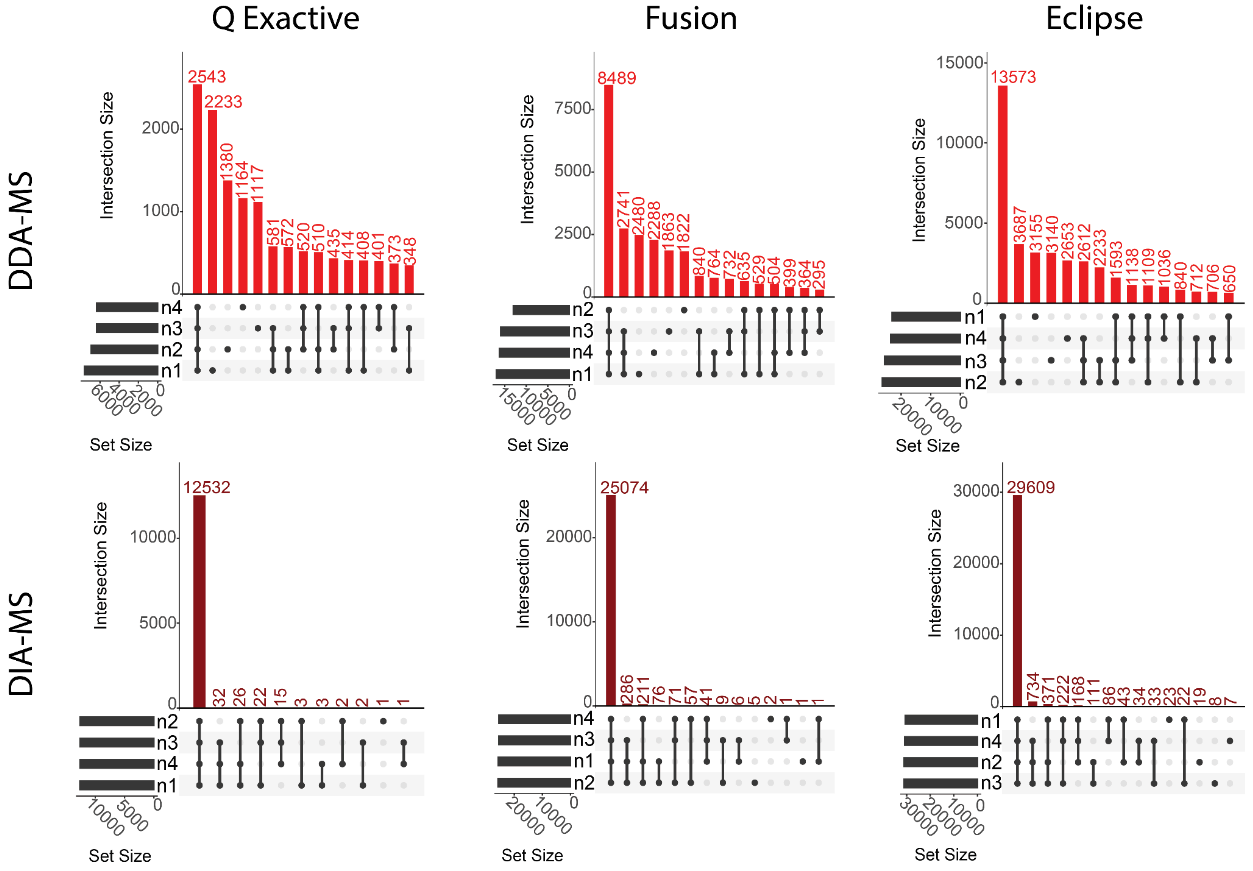

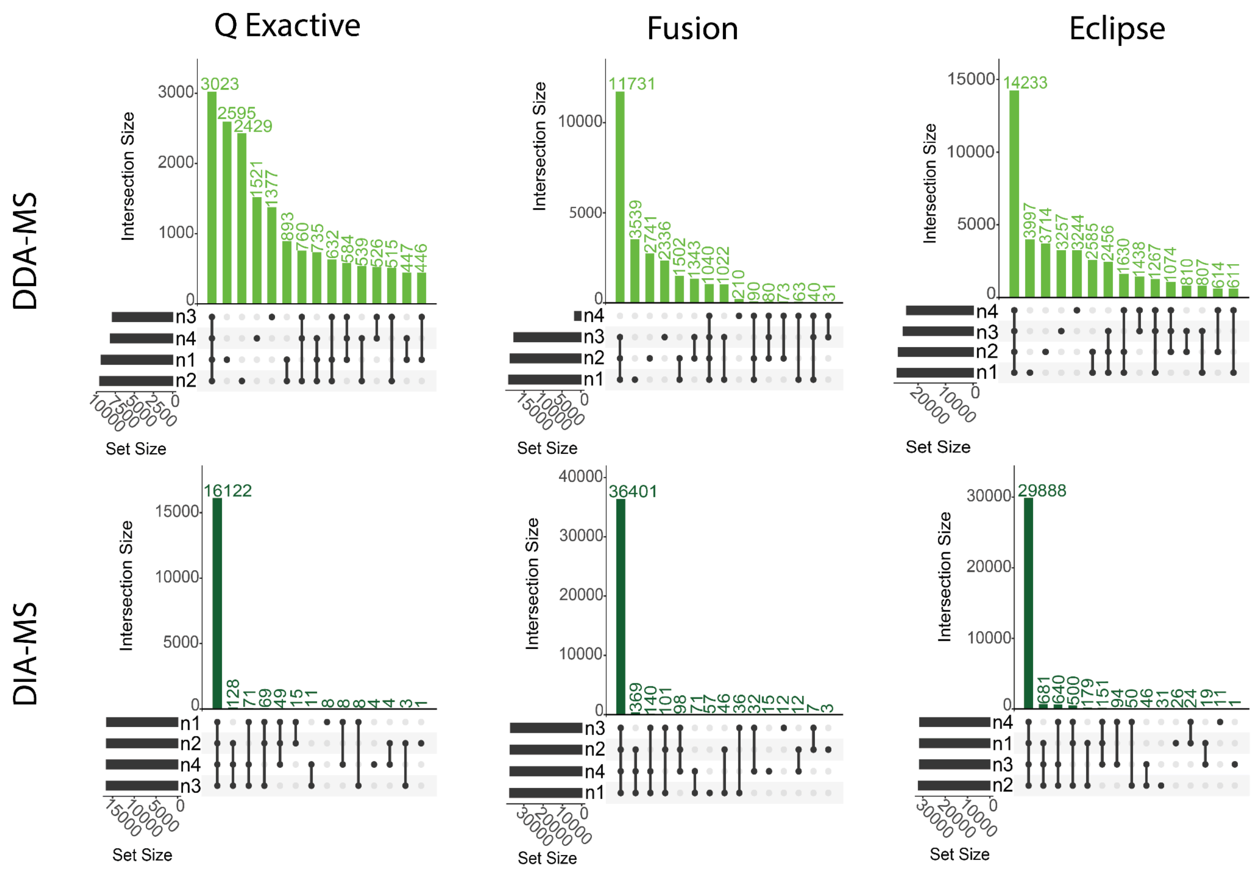

Q Exactive
Fusion
Eclipse
DDA-MS
DIA-MS
Intersection Size
Set Size

Supplemental Table S2: A) Proteins and B) peptides identified via DDA-MS and DIA-MS specific to the species within the microbiome

A)

| | QE Even Cell | | QE Even Protein | | QE Uneven | | Fusion Even Cell | | Fusion Even Protein | | Fusion Uneven | | Eclipse Even Cell | | Eclipse Even Protein | | Eclipse Uneven | |
|---|---|---|---|---|---|---|---|---|---|---|---|---|---|---|---|---|---|---|
| Species | DDA | DIA | DDA | DIA | DDA | DIA | DDA | DIA | DDA | DIA | DDA | DIA | DDA | DIA | DDA | DIA | DDA | DIA |
| 137 | 1 | 71 | 17 | 81 | 3 | 27 | 2 | 138 | 31 | 172 | 7 | 53 | 5 | 194 | 35 | 191 | 8 | 55 |
| 259 | 68 | 106 | 383 | 508 | 112 | 163 | 125 | 188 | 555 | 804 | 202 | 326 | 192 | 245 | 658 | 782 | 271 | 337 |
| 841 | 144 | 232 | 301 | 460 | 109 | 204 | 251 | 464 | 461 | 905 | 234 | 490 | 502 | 691 | 748 | 987 | 361 | 521 |
| AK199 | 124 | 66 | 141 | 83 | 41 | 23 | 197 | 128 | 230 | 149 | 100 | 62 | 388 | 179 | 375 | 156 | 134 | 61 |
| Am2 | 28 | 44 | 96 | 137 | 19 | 30 | 53 | 116 | 162 | 339 | 45 | 90 | 113 | 167 | 250 | 362 | 57 | 82 |
| ATN | 111 | 216 | 149 | 249 | 225 | 391 | 218 | 391 | 258 | 538 | 408 | 716 | 413 | 562 | 413 | 565 | 617 | 780 |
| BS | 194 | 267 | 126 | 190 | 22 | 48 | 325 | 493 | 205 | 397 | 54 | 125 | 506 | 639 | 307 | 416 | 91 | 130 |
| BXL | 10 | 13 | 8 | 17 | 14 | 27 | 23 | 32 | 23 | 39 | 34 | 47 | 33 | 43 | 34 | 50 | 51 | 45 |
| CRH | 451 | 673 | 48 | 91 | 63 | 108 | 718 | 1134 | 87 | 191 | 121 | 247 | 1118 | 1605 | 121 | 211 | 168 | 261 |
| Cup | 67 | 125 | 163 | 272 | 584 | 908 | 147 | 272 | 294 | 613 | 956 | 1490 | 292 | 426 | 456 | 656 | 1284 | 1575 |
| CV | 159 | 99 | 105 | 129 | 44 | 45 | 271 | 189 | 173 | 259 | 81 | 109 | 475 | 261 | 282 | 282 | 109 | 107 |
| DVH | 3 | 5 | 5 | 8 | 22 | 45 | 6 | 16 | 9 | 18 | 44 | 116 | 6 | 12 | 10 | 12 | 88 | 148 |
| ES18 | 1 | 2 | 0 | 3 | 1 | 0 | 2 | 2 | 2 | 2 | 0 | 0 | 2 | 1 | 2 | 2 | 0 | 0 |
| F0 | 0 | 0 | 2 | 3 | 1 | 1 | 0 | 2 | 3 | 5 | 1 | 2 | 1 | 0 | 4 | 4 | 1 | 2 |
| F2 | 1 | 1 | 1 | 1 | 0 | 0 | 1 | 1 | 1 | 1 | 1 | 1 | 1 | 1 | 1 | 1 | 1 | 0 |
| HB2 | 87 | 158 | 132 | 218 | 24 | 51 | 179 | 360 | 251 | 529 | 64 | 192 | 396 | 602 | 431 | 672 | 143 | 263 |
| K12 | 192 | 272 | 238 | 327 | 302 | 416 | 292 | 413 | 351 | 574 | 453 | 683 | 410 | 499 | 453 | 567 | 583 | 660 |
| KF7 | 130 | 194 | 111 | 170 | 30 | 54 | 215 | 358 | 163 | 356 | 67 | 126 | 420 | 566 | 289 | 407 | 108 | 129 |
| LT2 | 127 | 173 | 145 | 210 | 295 | 454 | 205 | 356 | 228 | 492 | 498 | 840 | 370 | 462 | 360 | 475 | 705 | 865 |
| M13 | 0 | 0 | 0 | 0 | 0 | 0 | 1 | 1 | 0 | 1 | 0 | 0 | 0 | 0 | 0 | 0 | 0 | 0 |
| Ne1 | 2 | 6 | 4 | 5 | 7 | 13 | 8 | 13 | 6 | 9 | 11 | 21 | 11 | 17 | 11 | 13 | 17 | 24 |
| Nm1 | 6 | 3 | 5 | 5 | 6 | 14 | 7 | 8 | 6 | 9 | 10 | 24 | 11 | 15 | 12 | 21 | 17 | 22 |
| Nu1 | 3 | 5 | 4 | 5 | 3 | 5 | 6 | 11 | 7 | 11 | 8 | 9 | 10 | 11 | 14 | 10 | 10 | 10 |
| NV | 3 | 10 | 83 | 161 | 10 | 23 | 9 | 16 | 176 | 357 | 20 | 75 | 18 | 37 | 250 | 371 | 47 | 88 |
| P22 | 0 | 0 | 2 | 2 | 1 | 1 | 0 | 0 | 1 | 4 | 1 | 1 | 0 | 0 | 3 | 6 | 2 | 2 |
| PaD | 41 | 252 | 100 | 317 | 15 | 92 | 83 | 519 | 182 | 687 | 44 | 241 | 157 | 733 | 254 | 699 | 77 | 245 |
| PD | 150 | 232 | 140 | 225 | 97 | 155 | 252 | 421 | 249 | 482 | 203 | 373 | 433 | 569 | 372 | 511 | 289 | 412 |
| Pfl | 103 | 168 | 119 | 217 | 230 | 364 | 181 | 331 | 222 | 507 | 419 | 697 | 316 | 433 | 321 | 452 | 602 | 733 |
| SMS | 145 | 214 | 173 | 245 | 345 | 538 | 224 | 404 | 255 | 556 | 592 | 987 | 425 | 599 | 417 | 615 | 827 | 1068 |

| | | | | | | | | | | | | | | | | | | |
|---|---|---|---|---|---|---|---|---|---|---|---|---|---|---|---|---|---|---|
| VF | 11 | 17 | 19 | 26 | 12 | 18 | 16 | 24 | 24 | 58 | 16 | 28 | 23 | 37 | 39 | 62 | 23 | 38 |

B)

| | QE Equal Cell | | QE Equal Protein | | QE Uneven | | Fusion Equal Cell | | Fusion Equal Protein | | Fusion Uneven | | Eclipse Equal Cell | | Eclipse Equal Protein | | Eclipse Uneven | |
|---|---|---|---|---|---|---|---|---|---|---|---|---|---|---|---|---|---|---|
| Species | DDA | DIA | DDA | DIA | DDA | DIA | DDA | DIA | DDA | DIA | DDA | DIA | DDA | DIA | DDA | DIA | DDA | DIA |
| 137 | 4 | 209 | 39 | 206 | 5 | 51 | 8 | 455 | 92 | 527 | 22 | 149 | 16 | 559 | 118 | 482 | 26 | 117 |
| 259 | 138 | 290 | 1470 | 2610 | 267 | 524 | 374 | 618 | 2741 | 4607 | 685 | 1350 | 597 | 700 | 3411 | 3829 | 1035 | 1183 |
| 841 | 334 | 641 | 732 | 1469 | 234 | 494 | 796 | 1427 | 1576 | 3485 | 667 | 1417 | 1469 | 1838 | 2387 | 2913 | 1095 | 1305 |
| AK199 | 274 | 141 | 337 | 174 | 82 | 33 | 590 | 318 | 719 | 386 | 260 | 120 | 1063 | 416 | 1082 | 336 | 388 | 118 |
| Am2 | 65 | 106 | 234 | 403 | 43 | 46 | 149 | 276 | 476 | 1081 | 143 | 187 | 316 | 333 | 780 | 897 | 187 | 150 |
| ATN | 223 | 492 | 315 | 630 | 472 | 1067 | 580 | 1066 | 731 | 1648 | 1233 | 2496 | 1094 | 1385 | 1143 | 1422 | 1965 | 2407 |
| BS | 548 | 1028 | 350 | 646 | 66 | 103 | 1249 | 2094 | 747 | 1610 | 192 | 321 | 1983 | 2380 | 1103 | 1315 | 308 | 307 |
| BXL | 44 | 25 | 56 | 24 | 60 | 33 | 155 | 55 | 173 | 69 | 191 | 60 | 178 | 69 | 192 | 65 | 214 | 51 |
| CRH | 1315 | 2383 | 209 | 206 | 235 | 262 | 2779 | 4434 | 520 | 494 | 676 | 663 | 4008 | 5178 | 623 | 412 | 853 | 624 |
| Cup | 141 | 275 | 380 | 751 | 1638 | 3461 | 428 | 708 | 900 | 2056 | 3694 | 6962 | 772 | 951 | 1367 | 1709 | 5546 | 6752 |
| CV | 396 | 256 | 261 | 357 | 97 | 85 | 847 | 490 | 526 | 761 | 237 | 240 | 1449 | 613 | 838 | 662 | 327 | 221 |
| DVH | 15 | 5 | 21 | 8 | 79 | 131 | 56 | 18 | 69 | 19 | 200 | 337 | 60 | 14 | 78 | 13 | 307 | 350 |
| ES18 | 1 | 2 | 1 | 4 | 1 | 0 | 2 | 5 | 2 | 6 | 1 | 0 | 4 | 3 | 3 | 3 | 1 | 0 |
| F0 | 1 | 0 | 3 | 8 | 1 | 1 | 2 | 2 | 10 | 19 | 2 | 4 | 1 | 0 | 14 | 14 | 3 | 3 |
| F2 | 1 | 2 | 1 | 1 | 0 | 0 | 2 | 2 | 2 | 2 | 1 | 2 | 2 | 2 | 2 | 1 | 84 | 0 |
| HB2 | 207 | 453 | 331 | 693 | 56 | 119 | 569 | 1257 | 790 | 2064 | 190 | 516 | 1234 | 1958 | 1417 | 2245 | 1006 | 651 |
| K12 | 522 | 951 | 703 | 1327 | 964 | 1746 | 1079 | 1659 | 1355 | 2473 | 1868 | 3153 | 1488 | 1799 | 1784 | 2051 | 2072 | 2976 |
| KF7 | 267 | 464 | 212 | 371 | 59 | 82 | 620 | 1000 | 454 | 948 | 188 | 238 | 1141 | 1318 | 787 | 862 | 771 | 213 |
| LT2 | 260 | 441 | 320 | 551 | 700 | 1242 | 540 | 942 | 629 | 1424 | 1467 | 2919 | 920 | 1091 | 957 | 1174 | 1769 | 2674 |
| M13 | 0 | 0 | 0 | 0 | 0 | 0 | 2 | 1 | 0 | 1 | 0 | 0 | 0 | 0 | 0 | 0 | 0 | 0 |
| Ne1 | 12 | 7 | 16 | 7 | 19 | 15 | 35 | 15 | 43 | 10 | 56 | 37 | 47 | 20 | 54 | 13 | 66 | 32 |
| Nm1 | 18 | 4 | 16 | 6 | 16 | 14 | 41 | 10 | 47 | 12 | 60 | 31 | 54 | 17 | 61 | 23 | 68 | 27 |
| Nu1 | 14 | 5 | 15 | 5 | 16 | 7 | 44 | 12 | 48 | 11 | 45 | 11 | 53 | 12 | 63 | 10 | 80 | 10 |
| NV | 23 | 26 | 194 | 437 | 41 | 54 | 64 | 41 | 505 | 1214 | 101 | 160 | 81 | 68 | 715 | 968 | 122 | 167 |
| P22 | 0 | 0 | 8 | 14 | 1 | 3 | 1 | 0 | 12 | 24 | 8 | 8 | 1 | 0 | 18 | 25 | 50 | 7 |
| PaD | 75 | 736 | 180 | 832 | 40 | 191 | 189 | 1636 | 401 | 2068 | 120 | 578 | 342 | 1948 | 552 | 1680 | 414 | 502 |
| PD | 326 | 598 | 313 | 607 | 201 | 357 | 707 | 1222 | 707 | 1455 | 541 | 1005 | 1256 | 1513 | 1089 | 1322 | 1124 | 1048 |
| Pfl | 243 | 418 | 290 | 629 | 569 | 1098 | 517 | 944 | 660 | 1554 | 1374 | 2536 | 892 | 1086 | 965 | 1196 | 2496 | 2441 |
| SMS | 360 | 709 | 449 | 825 | 1059 | 2127 | 783 | 1433 | 918 | 2142 | 2340 | 4524 | 1360 | 1759 | 1411 | 1830 | 2726 | 4384 |
| VF | 24 | 44 | 52 | 95 | 24 | 47 | 66 | 86 | 117 | 195 | 74 | 99 | 98 | 103 | 149 | 170 | 90 | 95 |

Supplemental Figure S2: Scatter plots of reference and measured percent abundances of species in the UNEVEN samples. Average Spearman correlation values show no significant difference to one another.

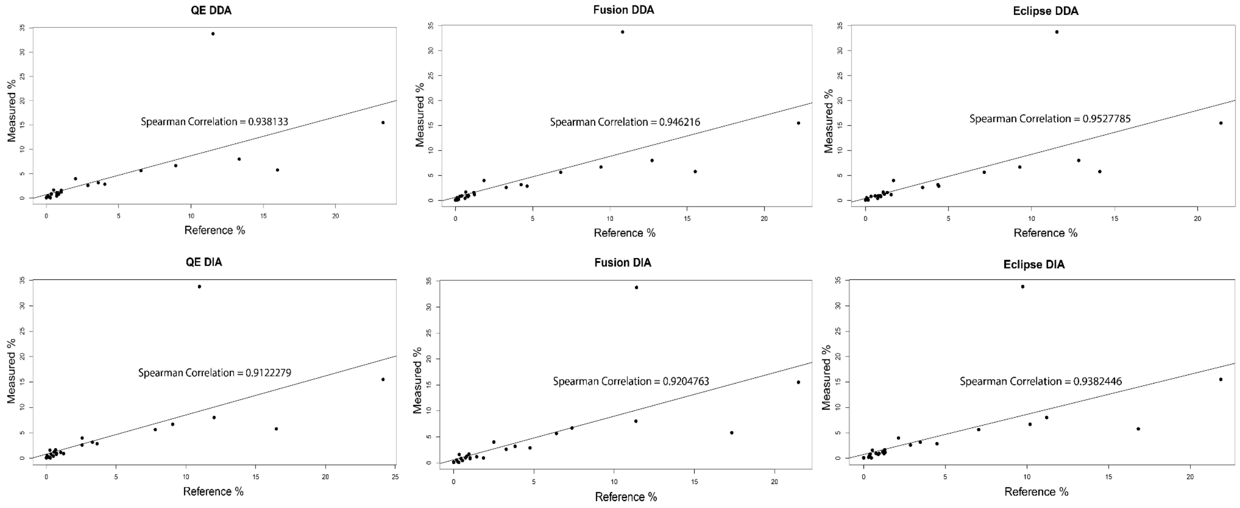

Supplemental Table S3: Fractional abundances of microbiome species in a) Equal Cell Number samples, b) Equal Protein Amount samples, and c) Uneven samples. Species in each sample type are listed from most to least abundant. DDA and DIA abundances for each species were compared via Student's t-test, with the p-value given indicating the significance of the difference between the values.

a)

| ECN | Q Exactive | | | Fusion | | | Eclipse | | |
|---|---|---|---|---|---|---|---|---|---|
| | DDA | DIA | p-value | DDA | DIA | p-value | DDA | DIA | p-value |
| CRH | 0.041598 | 0.047943 | 0.002366 | 0.072735 | 0.080826 | 0.108138 | 0.10462 | 0.114422 | 0.003014 |
| CV | 0.046806 | 0.024359 | 0.000152 | 0.081956 | 0.046559 | 9.39E-05 | 0.136532 | 0.064443 | 8.13E-07 |
| Pfl | 0.023745 | 0.028596 | 0.058204 | 0.042851 | 0.056298 | 0.009859 | 0.067447 | 0.073617 | 0.191603 |
| KF7 | 0.029227 | 0.034741 | 0.023494 | 0.052403 | 0.064147 | 0.009941 | 0.092613 | 0.101488 | 8.82E-05 |
| PD | 0.036185 | 0.046267 | 0.007819 | 0.063037 | 0.084049 | 0.000183 | 0.103863 | 0.113596 | 0.020628 |
| SMS | 0.04428 | 0.053625 | 0.03936 | 0.075828 | 0.101229 | 0.002935 | 0.128512 | 0.150339 | 0.003591 |
| AK199 | 0.041793 | 0.018177 | 1.55E-06 | 0.069058 | 0.035114 | 0.000161 | 0.127582 | 0.04916 | 9.89E-07 |
| HB2 | 0.049566 | 0.072065 | 0.001891 | 0.104728 | 0.16423 | 3.94E-05 | 0.206373 | 0.274897 | 9.47E-07 |
| ATN | 0.027246 | 0.040471 | 1.11E-06 | 0.053039 | 0.073298 | 0.000212 | 0.094917 | 0.105468 | 0.013085 |
| BS | 0.052195 | 0.063708 | 0.003601 | 0.09419 | 0.117633 | 2.59E-05 | 0.139048 | 0.15247 | 0.015094 |
| 841 | 0.027616 | 0.034763 | 7.65E-05 | 0.050853 | 0.069451 | 0.001759 | 0.092464 | 0.10339 | 0.000412 |
| LT2 | 0.038252 | 0.0448 | 0.007797 | 0.065807 | 0.092324 | 0.002154 | 0.112033 | 0.119684 | 0.003062 |
| K12 | 0.051999 | 0.064672 | 6.89E-05 | 0.082699 | 0.098346 | 0.00352 | 0.111851 | 0.118753 | 0.001998 |
| Cup | 0.015494 | 0.019763 | 0.013909 | 0.037826 | 0.042964 | 0.147164 | 0.06419 | 0.067273 | 0.238742 |
| Am2 | 0.011677 | 0.011355 | 0.790402 | 0.024323 | 0.029935 | 0.03558 | 0.048194 | 0.043097 | 0.199194 |
| PaD | 0.010355 | 0.044138 | 2.07E-08 | 0.022771 | 0.091128 | 2.67E-09 | 0.037996 | 0.128598 | 4.81E-11 |
| VF | 0.009589 | 0.011855 | 0.286738 | 0.022838 | 0.016736 | 0.001836 | 0.030335 | 0.025976 | 0.031126 |
| M13 | 0.027778 | 0 | 0.355918 | 0.166667 | 0.111111 | 0.133975 | 0 | 0 | - |
| BXL | 0.004823 | 0.00152 | 7.28E-06 | 0.016517 | 0.003742 | 0.000301 | 0.018475 | 0.004999 | 1.24E-05 |
| F2 | 0.1875 | 0.25 | 0.355918 | 0.25 | 0.25 | - | 0.25 | 0.25 | - |
| ES18 | 0.013158 | 0.026316 | 0.133975 | 0.026316 | 0.026316 | 1 | 0.039474 | 0.013158 | 0.002714 |
| 137 | 0.004428 | 0.114332 | 4.14E-11 | 0.010467 | 0.22182 | 2.98E-11 | 0.020531 | 0.312802 | 2.54E-10 |
| NV | 0.004256 | 0.003212 | 0.2914 | 0.014616 | 0.00514 | 0.000232 | 0.018712 | 0.011966 | 0.002995 |

| | | | | | | | | | |
|---|---|---|---|---|---|---|---|---|---|
| 259 | 0.032604 | 0.041022 | 0.00391 | 0.060275 | 0.072755 | 0.027848 | 0.088913 | 0.094814 | 0.000461 |
| F0 | 0.002273 | 0 | 0.133975 | 0.007955 | 0.007955 | 1 | 0.005682 | 0 | 0.094133 |
| P22 | 0.00463 | 0 | 0.355918 | 0.013889 | 0 | 0.168227 | 0.013889 | 0 | 0.168227 |
| DVH | 0.004226 | 0.001433 | 0.0005 | 0.015759 | 0.004585 | 3.97E-05 | 0.016547 | 0.003295 | 8.83E-05 |
| Ne1 | 0.005015 | 0.002561 | 0.013133 | 0.014405 | 0.005548 | 0.000177 | 0.019206 | 0.007149 | 1.42E-05 |
| Nm1 | 0.006285 | 0.001109 | 3.23E-07 | 0.014418 | 0.002957 | 7.45E-05 | 0.018762 | 0.00536 | 0.000221 |
| Nu1 | 0.004993 | 0.001816 | 0.008695 | 0.015704 | 0.003994 | 0.000197 | 0.019154 | 0.003903 | 4.78E-06 |

b)

| EPA | Q Exactive | | | Fusion | | | Eclipse | | |
|---|---|---|---|---|---|---|---|---|---|
| | DDA | DIA | p-value | DDA | DIA | p-value | DDA | DIA | p-value |
| LT2 | 0.005187 | 0.054461 | 2.24E-11 | 0.070799 | 0.127658 | 3.27E-08 | 0.110931 | 0.12312 | 0.027458 |
| PD | 0.082152 | 0.044919 | 5.48E-06 | 0.065033 | 0.096177 | 4.39E-06 | 0.093631 | 0.101917 | 0.103823 |
| BS | 0.084884 | 0.045216 | 0.000186 | 0.065378 | 0.094667 | 6.76E-06 | 0.092818 | 0.099141 | 0.183962 |
| PaD | 0.030932 | 0.055546 | 1.75E-06 | 0.041945 | 0.120612 | 7.27E-12 | 0.05616 | 0.122631 | 8.71E-08 |
| AK199 | 0.033737 | 0.022859 | 0.000463 | 0.07973 | 0.041036 | 8.74E-06 | 0.123933 | 0.043032 | 2.72E-05 |
| KF7 | 0.03371 | 0.030393 | 0.112695 | 0.043886 | 0.063789 | 0.000176 | 0.070109 | 0.072889 | 0.577765 |
| CV | 0.038789 | 0.031759 | 0.020839 | 0.059571 | 0.063764 | 0.000715 | 0.089973 | 0.069623 | 0.003549 |
| ATN | 0.010223 | 0.046614 | 4.99E-09 | 0.063356 | 0.100966 | 2.8E-07 | 0.095808 | 0.105937 | 0.02545 |
| SMS | 0.001317 | 0.061465 | 4.28E-15 | 0.083668 | 0.139488 | 3.63E-08 | 0.127321 | 0.154227 | 0.004756 |
| Cup | 0.025099 | 0.042964 | 3.1E-06 | 0.061581 | 0.096877 | 1.13E-06 | 0.093478 | 0.103755 | 0.01986 |
| Pfl | 0.037021 | 0.036894 | 0.945394 | 0.052596 | 0.086213 | 0.002038 | 0.073191 | 0.076894 | 0.670084 |
| BXL | 0.015669 | 0.001988 | 5.27E-08 | 0.019089 | 0.004531 | 4.5E-07 | 0.020989 | 0.005847 | 1.54E-07 |
| 137 | 0.032609 | 0.130435 | 2.94E-06 | 0.061997 | 0.27657 | 2.72E-11 | 0.073269 | 0.307568 | 5.44E-10 |
| 259 | 0.000484 | 0.196594 | 9.73E-19 | 0.236068 | 0.311146 | 8.89E-08 | 0.275348 | 0.302632 | 0.012693 |
| Am2 | 0.000516 | 0.035419 | 6.84E-13 | 0.058903 | 0.087548 | 3.89E-07 | 0.085935 | 0.09329 | 0.181881 |
| K12 | 0.010531 | 0.077701 | 0.000618 | 0.097394 | 0.136602 | 2.5E-06 | 0.125595 | 0.159864 | 7.92E-05 |
| HB2 | 0.088282 | 0.099703 | 0.477159 | 0.138648 | 0.241663 | 3.73E-08 | 0.223732 | 0.258908 | 0.002279 |
| CRH | 0.016433 | 0.006505 | 0.002568 | 0.028249 | 0.013599 | 1.97E-08 | 0.032348 | 0.015025 | 1.11E-06 |
| 841 | 0.022938 | 0.068889 | 2.23E-06 | 0.086477 | 0.135496 | 1.45E-07 | 0.132727 | 0.14777 | 0.013839 |
| VF | 0.089435 | 0.018131 | 0.034949 | 0.03016 | 0.040098 | 8.55E-07 | 0.043236 | 0.042887 | 0.938328 |
| NV | 0.005381 | 0.051558 | 4.23E-10 | 0.075169 | 0.114761 | 0.000288 | 0.097334 | 0.119017 | 0.033991 |
| M13 | 0 | 0 | - | 0.027778 | 0.111111 | 0.024008 | 0 | 0 | - |
| F2 | 0.25 | 0.25 | - | 0.25 | 0.25 | - | 0.25 | 0.25 | - |
| P22 | 0.037037 | 0.032407 | 0.62022 | 0.050926 | 0.074074 | 0.094133 | 0.083333 | 0.111111 | 0.168227 |
| F0 | 0.010227 | 0.013636 | 0.168227 | 0.020455 | 0.022727 | 0.5847 | 0.029545 | 0.018182 | 0.027811 |

| | | | | | | | | | |
|---|---|---|---|---|---|---|---|---|---|
| ES18 | 0.009868 | 0.039474 | 0.000105 | 0.029605 | 0.026316 | 0.355918 | 0.029605 | 0.026316 | 0.355918 |
| DVH | 0.045129 | 0.002292 | 6.69E-05 | 0.019413 | 0.005158 | 2.36E-06 | 0.021848 | 0.003438 | 1.74E-06 |
| Ne1 | 0.074584 | 0.002027 | 0.000257 | 0.017926 | 0.003948 | 2.4E-05 | 0.021874 | 0.005335 | 2.41E-07 |
| Nm1 | 0.061738 | 0.001848 | 0.014631 | 0.016913 | 0.003235 | 3.67E-06 | 0.021257 | 0.007671 | 8.6E-06 |
| Nu1 | 0.006445 | 0.001816 | 0.088498 | 0.017248 | 0.003994 | 4.74E-05 | 0.022331 | 0.00345 | 6.96E-06 |

c)

| Uneven | Q Exactive | | | Fusion | | | Eclipse | | |
|---|---|---|---|---|---|---|---|---|---|
| | DDA | DIA | p-value | DDA | DIA | p-value | DDA | DIA | p-value |
| LT2 | 0.083182 | 0.117674 | 0.0009 | 0.1427 | 0.217713 | 1.29E-05 | 0.200402 | 0.224196 | 0.049676 |
| Cup | 0.102964 | 0.143478 | 3.97E-07 | 0.169763 | 0.235494 | 3.39E-05 | 0.227708 | 0.248972 | 0.002486 |
| SMS | 0.098595 | 0.135035 | 9.65E-09 | 0.167963 | 0.247679 | 2.96E-05 | 0.234069 | 0.267875 | 0.000163 |
| Pfl | 0.047362 | 0.061957 | 3.29E-06 | 0.08766 | 0.118553 | 2.13E-05 | 0.122723 | 0.124766 | 0.236677 |
| K12 | 0.079069 | 0.09906 | 0.003002 | 0.120895 | 0.162542 | 6.22E-06 | 0.154569 | 0.157009 | 0.63211 |
| ATN | 0.049709 | 0.073345 | 2.51E-05 | 0.093744 | 0.134356 | 3.23E-05 | 0.136278 | 0.146361 | 0.007081 |
| CRH | 0.012797 | 0.007682 | 0.00041 | 0.034576 | 0.017609 | 9.49E-05 | 0.03978 | 0.018625 | 1.4E-10 |
| PD | 0.025105 | 0.030994 | 0.036784 | 0.0556 | 0.074516 | 0.002524 | 0.077361 | 0.082252 | 0.385001 |
| VF | 0.011681 | 0.012378 | 0.346422 | 0.026848 | 0.019526 | 0.01308 | 0.031206 | 0.026151 | 0.071268 |
| HB2 | 0.015418 | 0.023298 | 0.001473 | 0.046368 | 0.087597 | 2.25E-06 | 0.087825 | 0.12026 | 0.001514 |
| 259 | 0.050019 | 0.062984 | 0.001116 | 0.09404 | 0.126064 | 6.83E-05 | 0.121904 | 0.130418 | 0.170644 |
| AK199 | 0.016249 | 0.006265 | 0.001151 | 0.045098 | 0.016937 | 2.68E-06 | 0.056803 | 0.016662 | 4.33E-05 |
| CV | 0.01665 | 0.0111 | 5.33E-05 | 0.038234 | 0.026764 | 9.34E-05 | 0.046929 | 0.026394 | 9.75E-06 |
| KF7 | 0.009638 | 0.009638 | 1 | 0.026986 | 0.022548 | 0.017457 | 0.036624 | 0.023041 | 6.01E-05 |
| 137 | 0.006441 | 0.043478 | 5.62E-08 | 0.026973 | 0.085749 | 8.7E-08 | 0.025765 | 0.088164 | 3.66E-08 |
| Am2 | 0.009355 | 0.007677 | 0.021707 | 0.029548 | 0.023097 | 0.002101 | 0.033484 | 0.021161 | 0.000666 |
| DVH | 0.012249 | 0.012894 | 0.520897 | 0.029656 | 0.033166 | 0.004196 | 0.044986 | 0.042407 | 0.157125 |
| PaD | 0.005923 | 0.016058 | 3.78E-07 | 0.017111 | 0.042296 | 2.06E-07 | 0.022552 | 0.042954 | 3.7E-09 |
| 841 | 0.021741 | 0.03046 | 1.48E-05 | 0.050142 | 0.073342 | 6.33E-05 | 0.073043 | 0.077982 | 0.024008 |
| NV | 0.007549 | 0.007388 | 0.873684 | 0.021924 | 0.024012 | 0.083052 | 0.031561 | 0.028188 | 0.094264 |
| BS | 0.009962 | 0.011393 | 0.04925 | 0.027977 | 0.029707 | 0.007255 | 0.038654 | 0.030959 | 2.47E-05 |
| Nu1 | 0.005628 | 0.001816 | 1.91E-05 | 0.01634 | 0.003359 | 4.32E-07 | 0.022059 | 0.00354 | 0.000225 |
| BXL | 0.007104 | 0.003128 | 4.06E-06 | 0.02134 | 0.005437 | 9.62E-06 | 0.023913 | 0.005203 | 1.39E-08 |
| Nm1 | 0.0061 | 0.005176 | 0.421936 | 0.02098 | 0.008965 | 2.27E-05 | 0.023383 | 0.007948 | 0.000282 |
| M13 | 0.027778 | 0 | 0.355918 | 0 | 0 | - | 0 | 0 | - |

| | | | | | | | | | |
|---|---|---|---|---|---|---|---|---|---|
| P22 | 0.013889 | 0.018519 | 0.355918 | 0.060185 | 0.018519 | 0.003326 | 0.041667 | 0.037037 | 0.355918 |
| F0 | 0.003409 | 0.004545 | 0.355918 | 0.007955 | 0.009091 | 0.779559 | 0.010227 | 0.009091 | 0.355918 |
| ES18 | 0.006579 | 0 | 0.133975 | 0.016447 | 0 | 0.14663 | 0.006579 | 0 | 0.355918 |
| F2 | 0 | 0 | - | 0.125 | 0.25 | 0.133975 | 0.1875 | 0 | 0.024008 |
| Ne1 | 0.007469 | 0.005442 | 0.000472 | 0.022514 | 0.008856 | 2.33E-05 | 0.026248 | 0.01035 | 5.92E-06 |

Supplemental Figure S3: Distribution of misidentified proteins matching to each entrapment genome. (A) A stacked bar chart depicting the average number of proteins identified with at least 1 unique peptide from each of the entrapment genomes added to the A1, A1I1, and A1I2 databases using the DIA method. (B) A stacked bar chart depicting the average number of proteins identified with at least 1 unique peptide from each of the entrapment genomes added to the A1, A1I1, and A1I2 databases using the DDA method

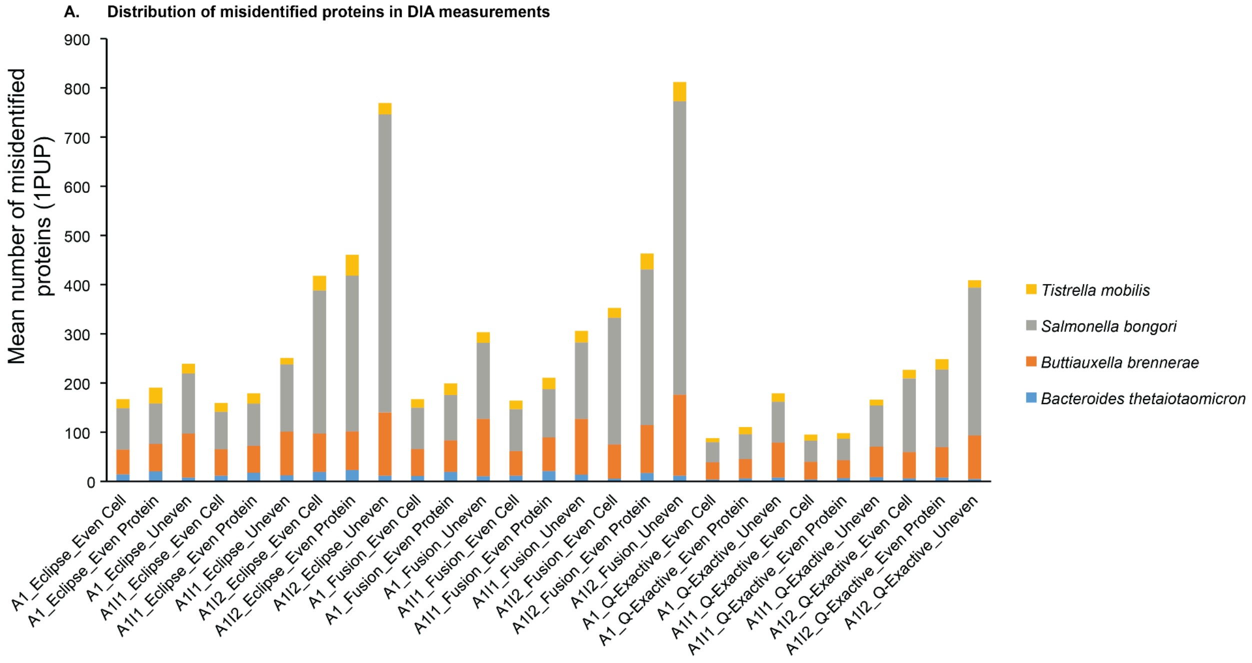


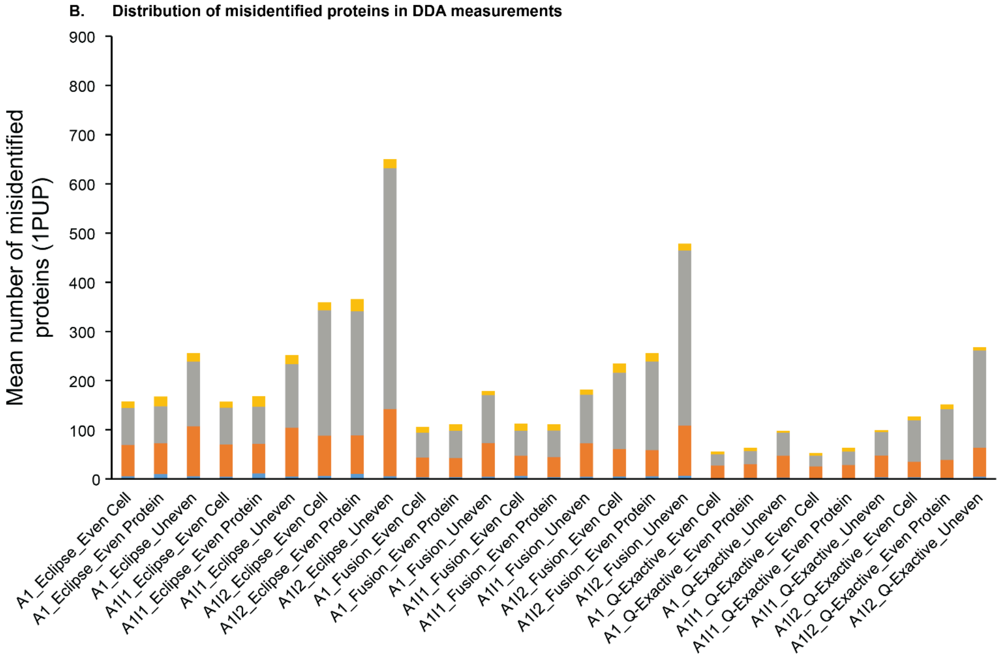
